# Proopiomelanocortin (POMC) is a negative regulator of tadpole aggression through opioid receptor signaling

**DOI:** 10.1101/2022.11.28.518266

**Authors:** Jordan E. McKinney, Sarah C. Ludington, Julie M. Butler, Lauren A. O’Connell

## Abstract

Aggression is a common behavioral response to limited environmental resources. Most research on the neural basis of aggression in vertebrates focuses on adult males, where sex steroid hormones and the ventromedial hypothalamus are important regulators of aggressive behavior. However, the young of many species also display aggression, although the neural basis of juvenile aggression is not well understood. Here we examine juvenile aggression in Mimic poison frog (*Ranitomeya imitator*) tadpoles, who live in small isolated pools with limited resources and display aggressive behaviors towards intruder tadpoles. We first conducted a longitudinal study of dyadic behavior and found aggressive behavior increases with tadpole age. We next evaluated which brain regions may be important for tadpole aggression by quantifying pS6-positive cells as a proxy for neural activity. We found more pS6-positive cells in the ventral hypothalamus of aggressive tadpoles compared to controls, similar to findings in adult mammals. We then quantified colocalization of pS6 with vasopressin and oxytocin cells and found no difference across behavior groups. Based on this information, we hypothesized that similar brain regions, but different neuromodulators, may promote aggression in juvenile tadpoles compared to the literature in adult animals. We then used an untargeted approach to molecularly profile pS6-positive neurons and found enrichment of the proopiomelanocortin (POMC) gene in aggressive tadpoles. As POMC is cleaved into several signaling peptides, we used pharmacology to target each pathway and discovered that blocking opioid receptors increases aggressive behavior. Together, this work suggests that POMC-derived B-endorphin is a negative regulator of juvenile aggression through the opioid receptor signaling. More broadly, this work suggests that similar brain regions, but different signaling molecules may be used to regulate aggression in adult and juvenile animals.

## Introduction

Aggression is a widespread behavior across the animal kingdom as an effective but risky social decision to defend access to resources. Most of what is known about the neural basis of aggression comes from studies in adult laboratory rodents. Decades of research in this area have highlighted important roles for sex steroid hormones in the elicitation and progression of aggression in mammals [1], birds [2,3], lizards [4,5], and fish [6,7]. Studies on the neural basis of aggression point to the ventromedial hypothalamus and adjacent brain areas, such as the medial amygdala, as regulating aggression in a wide variety of species, including adult rodents [8–11], reptiles [12], and birds [2,13]. Systems neuroscience research has linked these two observations together, showing that estrogen and progestin-sensitive neurons in the VMH regulate aggression in rodents [14–16]. However, most of these studies have been conducted in adult animals, where sex steroid hormones act in the brain to alter neural circuit dynamics to promote or inhibit aggression. In contrast, sex hormones are not present in high amounts in young, pre-pubertal animals, leaving a critical gap in our understanding of how aggression is regulated in the juvenile brain. This gap is important because the young of many species display aggression, including in birds [17], amphibians [18,19], and some mammals [20–22], sometimes with deadly consequences. Therefore, it is important to understand the neural basis of aggression in young and define the similarities and differences in mechanisms compared to adults.

In addition to sex steroid hormones, the hypothalamic-pituitary-adrenal (HPA) axis is important for regulating the development, elicitation, and enhancement of aggressive behavior in vertebrate animals [23–26], including humans [27,28]. The HPA axis regulates the release of corticotropin-releasing hormone and arginine vasopressin from the paraventricular nucleus of the hypothalamus, which stimulates secretion of adrenocorticotropic hormone (ACTH), a product of proopiomelanocortin (POMC), from the pituitary. Both vasopressin and corticotropin-releasing hormone have been linked to aggression in adult males [29,30]. The POMC peptide is cleaved into three smaller peptides: (1) alpha melanocyte-stimulating hormone (aMSH) that regulates aggression and appetite [31] (2) B-endorphins that are involved in hedonia and pain management through binding to opioid receptors [32], and (3) adrenocorticotropic hormone (ACTH) that is released from the pituitary and stimulates the production of cortisol in the adrenal gland [33]. The role of the HPA axis in regulating aggression in juveniles is not well understood, but is a promising site of regulation given the relative lack of sex steroid hormones levels compared to adults.

Little is known about the neural mechanisms of juvenile aggression as compared to adults, as few laboratory research organisms are naturally aggressive when young. Some laboratory rodent juveniles display “play fighting” activities [34], and the similarities and differences in mechanism compared to the offensive aggression seen in adult interactions is unclear. Play fighting in mouse juveniles does not include hard bites and is often in amicable context [34]. However, there are similarities when comparing juvenile play to adult aggression, such as the ventromedial hypothalamus being more active after play behavior [35]. The HPA axis can also regulate the switch to aggressive behavior from play fighting in other rodent species during development because of changes in glucocorticoid levels [36]. Additionally, nonapeptides like vasopressin and oxytocin have been proposed to regulate juvenile play behavior [37], similar to adult aggression in mammals and other vertebrates [38–41]. Despite this progress in understanding rodent play behavior, we still lack a circuit level understanding of juvenile aggression and whether similar or different brain regions and cell types govern aggression in juveniles as compared to adults. What is needed is a research organism where juveniles display robust offensive aggression in an ecologically relevant context.

Some species of amphibian tadpoles show aggressive, cannibalistic behavior during development. For example, wood frog tadpoles (*Lithobates sylvaticus*) display cannibalism based on resource competition [42]. Poison frog species (Family Dendrobatidae) also show variation in aggressive behavior that is linked to ecological resources in their environment [43]. Parents place tadpoles either in large pools of water where tadpoles are generally gregarious or in small pools contained in plants where tadpoles have evolved aggressive behavior, presumably in defense of their limited resources. For example, the Mimic poison frog (*Ranitomeya imitator*) and the Strawberry poison frog (*Oophaga pumilio*) place tadpoles into small pools contained within plants. These tadpoles have independently evolved begging behavior to solicit food from their parents and aggressive behavior in response to other tadpoles. Although this behavior is hypothesized to be linked to proximate resource limitations, aggressive behavior in *O. pumilio* tadpoles is unlinked to food deprivation [44], suggesting nutrition state and aggression are decoupled. We have previously demonstrated in tadpoles of a closely related species (*Dendrobates tinctorius*) that winners of aggressive encounters had increased neural activity in preoptic area nonapeptide neurons compared to losers, emphasizing the importance of the hypothalamus in tadpole aggression [18], similar to studies in adult vertebrates [27,45,46]. However, it is still unclear which brain regions or cell types influence aggression in juvenile tadpoles.

Here, we tested the hypothesis that similar brain regions, but different neuromodulators, would be involved in regulating aggression in juvenile tadpoles compared to the literature in adult animals. Specifically, we predicted that the ventral hypothalamus and amygdala would be more active in aggressive tadpoles and that we would be able to identify neuromodulators important for aggression that are not related to sex steroid hormones. We first examined aggressive behavior across development and mapped the neural distribution of active neurons across the brain during aggression as compared to control tadpoles. We then molecularly profiled active neurons to identify candidate neuromodulators of aggressive behavior and followed up on promising candidates using pharmacology. Our results highlight both conserved and novel neuromodulatory patterns regulating juvenile aggression.

## Methods

### Animals

*Ranitomeya imitator* tadpoles were bred from our laboratory colony at Stanford University. Breeding pairs are housed in 10 × 10 × 12 cm terrarium containing sphagnum moss, driftwood, live philodendron plants, and canisters for egg laying and tadpole deposition. Tadpoles were moved from the terraria to individual perforated cups in a large water tank maintained at 25oC. Tadpoles were exposed to shared water parameters with constant circulation and fed brine shrimp flakes and standard tadpole food pellets (Josh’s Frogs, Owosso, MI, USA) three times weekly. Each tadpole container contained sphagnum moss and tea leaves for additional nutrients. All procedures were approved by the Stanford University Animal Care and Use Committee (Protocol #33097).

### Behavior

For all aggression trials, tadpoles were placed in a clear arena (5 × 5 × 5 cm) with 50 mL of reverse osmosis conditioned (Josh’s Frogs Dechlorinator Tap Water Conditioner) DI water. Each arena was placed on a square LED light pad (10 × 10 × 15 cm) with mirrors placed on adjacent sides angled at 60o to allow simultaneous filming of the top, front, and side views. Each tadpole was filmed using GoPro cameras (GoPro HERO7 Black, 1080p, 240 fps; GoPro, San Mateo, CA, USA) placed above the arena. All behavior trials were conducted between 9:00 AM and 12:00 PM. Videos were scored using BORIS software [47]. The behaviors scored were the number of bites, number of bites with thrashing action, and number of chasing actions.

For the development study, one week old focal tadpoles (N = 10) (Gosner stage 25-26) were tested once per week in an aggression trial over a period of six weeks. Tadpole size and mass were recorded once a week at the beginning of the week (Supplemental Figure 1A). Focal tadpoles were consistently slightly larger than stimulus tadpoles, although this was not statistically significant (Table S1). The tadpoles were allowed to acclimate for 10 minutes in the arena before a smaller conspecific tadpole was placed in the tank for 30 minutes. Then, both tadpoles were removed and placed back in their home canisters.

For aggression trials for downstream neural studies, tadpoles were randomly assigned at Gosner stage 26-30 (no substantial limb development) into one of two experimental groups: exposed to a novel object stimulus (metal bolt control, N = 15) or exposed to a smaller tadpole (aggression, N = 15). Focal tadpoles were slightly larger than stimulus tadpoles (aggressive tadpole ave = 23.33mm, sd = 1.67mm; stimulus tadpole ave = 18.85mm, sd = 1.57mm; size difference ave = 1.24mm, sd = 1.57mm), although these differences were not significant (T(4)= 1.5919, p = 0.1866). Each tadpole acclimated for 10 minutes in the arena, after which the stimulus tadpole was added and behavior was recorded. After 30 minutes, stimuli were removed from the arena and focal tadpoles were placed in the dark for 15 minutes. Focal tadpoles were then anesthetized with topical 20% benzocaine, weighed and measured, and sacrificed by rapid decapitation.

### Immunohistochemistry

Whole tadpole heads were fixed with 4% paraformaldehyde (PFA) in 1X phosphate buffered saline (PBS) at 4oC overnight, rinsed in 1X PBS, and transferred to a 30% sucrose solution for cryoprotection at 4oC overnight. Tadpole heads were then embedded in mounting media (Tissue-Tek® O.C.T. Compound, Electron Microscopy Sciences, Hatfield, PA, USA), rapidly frozen on dry ice, and stored at -80oC until cryosectioning. Heads were sectioned at 15μm into three series on a cryostat. Sections were thaw-mounted onto SuperFrost Plus microscope slides (VWR International, Randor, PA, USA), allowed to dry completely, and then stored at -80oC.

We first examined nonapeptide cells, given their role in juvenile play behavior in mammals [48] and aggression in adult vertebrates [29]. We used double-label fluorescence immunohistochemistry to detect phosphorylated ribosomes (pS6, phospho-S6) as a proxy of neuronal activity [49] and either vasopressin or oxytocin peptides. Briefly, slides were removed from the -80oC and air-dried before being fixed in chilled 4% PFA in 1X PBS. Slides were blocked with a blocking solution (1X PBS, 0.2% BSA, 0.2% Triton-X, 5% NGS) at room temperature for 1 hour. Slides were then incubated in a mix of antibodies to detect pS6 (rabbit anti-pS6 (Invitrogen, cat #44-923G) at 1:500, validated in poison frogs previously [50]), and either vasopressin (pre-incubated with mesotocin, (mouse PS45, a gift from Hal Grainger)), or oxytocin (1:5000, MAB5296; Millipore Sigma, Burlington, MA, USA; validated in poison frogs previously [18]). After several washes in 1X PBS, slides were incubated in a mix of fluorescent secondary antibodies (1:200 AlexaFluor 488 anti-rabbit and AlexaFluor 594 anti-mouse in blocking solution) for two hours. After rinsing in water, all slides were cover slipped using Vectashield Hardset Mounting Medium with DAPI (Vector Laboratories, Burlingame, CA, USA) and stored at 4oC until imaging.

### Molecular profiling of brain cells using phosphoTRAP

After aggression assays, the tadpoles that engaged in aggressive behavior (N = 6) and tadpoles exposed to a control stimulus (N = 5, metal bolt) were collected. Tadpoles were euthanized as described above and whole brains were removed, cut into three parts representing the telencephalon, diencephalon, and hindbrain and placed in homogenization buffer (10mM HEPES solution pH 7.4, Fisher Scientific, #NC0689154), 150mM KCl solution (Sigma Aldrich, #44675), 5mM MgCl2 solution (Sigma Aldrich, #M2393) in water with additional additives (0.5mM DTT, Fisher Scientific, #FERR0861), protease inhibitors (Sigma-Aldrich, #11873580001), RNAsin (Fisher Scientific, #PRN2615), 100 ug/mL cycloheximide (Fisher Scientific, #50-200-8999), phosphatase inhibitor cocktail (Sigma-Aldrich, #P0044), and calcyculin (Sigma Aldrich, #C5552) in a 2 mL BeadBug™ homogenization tube (Sigma-Aldrich, St Louis, MO, USA). After homogenization, lysate was spun, transferred to a new microfuge tube, and clarified using 70 uL of 10% NP40 and 70 uL of DHPC per 1.0 mL of supernatant. Supernatant was spun at Rmax for 10 minutes and transferred to a new tube. A small aliquot of supernatant was stored at -80oC, representing INPUT RNA. The clarified supernatant was added to pS6 antibody-loaded beads (Protein A Dynabeads, Invitrogen) and incubated for 10 minutes on an end over end mixer. Beads were placed on a magnet and supernatant was removed. Beads were resuspended with 0.9 mL of cold 0.35M KCl IP wash buffer solution (5 mL of 1M HEPES, 87.5 mL of 2M KCl solution, 2.5 mL of 1M MgCl2, 50 mL of 10% NP40, 355mL milliQ water) with additives (5 μL of 1M DTT solution, 10 μL RNAsin, 5 μL of 100 mg/mL Cycloheximide, 2μL 1000X Calcyculin) per every 4mL needed and resuspended. This was repeated for a total of 4 washes, and then 350 uL of buffer RLT (Qiagen RNeasy Micro kit, #74034) was added and tubes were placed on ice for 5 minutes and then put back on the magnet. Supernatant containing immunoprecipitated (IP) RNA was transferred to a new tube. Both INPUT and IP RNA was purified using the RNeasy Micro kit (Qiagen, #74034) and stored at -80oC. Libraries were prepared for both the INPUT and IP RNA using the NEBNext Single Cell/Low Input RNA Library Prep Kit for Illumina) (NEB, #E6420S) following the manufacturer’s protocol. Libraries were pooled in equimolar amounts and sequenced on an Illumina HiSeq 2500.

### In situ hybridization

In situ hybridization was used to detect proopiomelanocortin (*pomc)*-expressing cells. Probes were created by PCR amplification of *pomc* from tadpole cDNA using the following primers: forward primer (sense): 5’-GGCAGCGACGGCAATAATA-3’, reverse primer (antisense): 5’-CCGTCGCTGCCGTTATTAT-3’ with a T3 polymerase recognition sequence (5’-AATTAACCCTCACTAAAGGG-3’) on the 5’ end; Integrated DNA Technologies (IDT, San Diego, CA, USA). The PCR products were confirmed on agarose gel at the appropriate size and only a single band was observed. PCR products were purified (MinElute PCR kit – Qiagen #28004) and used as template in a transcription reaction to incorporate DIG-labeled nucleotides into an RNA probe (Thermo Fisher MEGAscript T3 Transcription Reaction Kit). Probes were purified with G50 microcolumns (GE Illustra ProbeQuant G-50 microcolumns), confirmed on agarose gel, and then diluted 1:5 by adding 100 uL of pre-hybridization buffer (23mL MilliQ water, 50mL 100% Formamide, 25mL 20X SSC, 100uL Tween-20, 0.1g CHAPS, 1.0mL pH 8.0 0.5 M EDTA, 1.0mL 50mg/mL Torula RNA). To verify probe specificity, alternate series of sectioned tadpole brains were stained with the antisense and sense probes. No staining was visible using the sense probe (Figure S1). The purified PCR product was Sanger sequenced (GeneWiz, Inc, South San Francisco, CA, USA) using the primer for generating the probe to confirm the *pomc* sequence.

Slides were washed 3 times with 1X PBS for 5 minutes each, and then fixed with 4% PFA for 20 minutes. Slides were washed two times with 1X PBS for 5 minutes each and then incubated in proteinase K (PK) solution (10 ml proteinase K buffer (Thermo Fisher-# EO0491) and 5 μl of 20 mg/ml proteinase K) for 10 minutes. Then, slides were washed once with 1X PBS for 10 minutes, fixed with 4% PFA for 15 minutes, washed two times with 1X PBS for 5 minutes each, washed with milliQ water, and incubated with 0.1M triethanolamine-HCl (pH 8.0) containing 0.25% acetic anhydride to 1:400 for 10 minutes. Slides were washed with 1X PBS for 5 minutes and incubated in pre-warmed hybridization buffer for 2 hours at 60oC inside a covered hybridization chamber (small plastic container with paper towels on the bottom soaked in 1:1 formamide & milliQ water, egg-crate platform on top). The chamber was removed from the oven and the buffer removed. Next, 200 uL of the probe solution (1 mL hybridization buffer, 8 μL DIG-labeled *pomc* probe) was added to each slide and a HybriSlip hybridization coverslip (Invitrogen Hybrislip Hybridization cover, RNase free) was placed over the side. The lid was placed on the chamber and incubated at 60oC overnight.

The next morning, HybriSlips were removed from the slides by immersing the slides in pre-warmed 2X saline-sodium citrate/tween (SSC/Tw):50% Formamide and moving the slide back and forth. Slides were then washed two times in pre-warmed 2X SSC/Tw:50% Formamide for 40 minutes at room temperature and then 20 minutes at 60°C. Slides were then washed in pre-warmed 1:1 2X SSC/Tw:maleate buffer containing Tween-20 (MABT) for 15 min each at 60oC. Slides were then washed in pre-warmed MABT for 10 min each at 60oC. Slides were then transferred to a flat glass staining plate, and washed two times with MABT for 10 min each at room temperature. Slides were blocked with MABT containing 2% bovine serum albumin (BSA) for 3 hours at room temperature. Slides were incubated with anti-DIG antibody (Roche, #11093274910) diluted 1:5000 in blocking solution (MABT + 2% BSA) at 4oC overnight in a flat, humidified chamber.

The next morning, slides were washed with MABT three times for 20 minutes each at room temperature. The DIG probe was then visualized with Fast Red TR (Fast Red TR Salt 1,5-naphthalenedisulfonate salt, Sigma-Aldrich) by applying the solution to slides and incubating at 37oC inside a dark, humidified chamber. The reaction was stopped after two hours, when there was positive staining of cells in the area of interest and low background. Slides were then washed with 1X PBS once for 5 minutes at room temperature, and then three times for 10 minutes each. Slides were incubated with pS6 antibody solution (7.5mL blocking solution: 10 mL PBS, 1 mL NGS, 0.02 g BSA, 30 uL Triton-X, 2 uL pS6 antibody) overnight at 4oC in a humidified chamber. The next morning, slides were processed for immunohistochemical detection of pS6 (1:400 AlexaFluor 488 secondary antibody) as described above.

### Fluorescence microscopy and cell counting

Brain sections were imaged on a Leica compound light microscope connected to a QImaging Retiga 2000R camera and a fluorescent light source. Each section was visualized at three fluorescent wavelengths (594, 488, and 358 nm) and images were pseudo-colored to reflect cyan (pS6), pink (vasopressin, oxytocin, or *pomc*), or blue (DAPI, stains DNA). Brain regions containing POMC were identified using DAPI-stained nuclei while referencing a poison frog brain atlas [51]. For each brain region counted, FIJI software [52] was used to measure the area of each brain region before the number of neuropeptide-positive cells, pS6-positive cells, and colocalized cells were quantified within a single region for each section using the “Cell Counter” function. pS6 cells were counted in anterior amygdala (AA), bed nucleus of the stria terminalis (BST), lateral amygdala (LA), medial amygdala (MeA), medial pallium (MP), preoptic area of the hypothalamus (POA), and ventral hypothalamus (VH) because these areas are involved in the HPA-axis, which is implicated in aggressive behavior [27]. For nonapeptides, cells were quantified in the POA. For *pomc*, cells were quantified in the arcuate nucleus, POA, and VH.

### POMC pharmacology

Tadpoles were anesthetized in 0.01% MS-222 in frog water buffered in sodium bicarbonate for four minutes, and then injected using pulled glass micropipettes. Tadpoles were injected intracranially (icv, in the ventricle) or intraperitoneally (ip, in the body cavity) with 50.6 nl of a double-blinded drug, with varying concentration based on the drug type being tested. Tadpoles were weighed and measured before being placed back in their home tank for recovery. For most drug treatments, tadpoles were injected between 7:30-8:30 AM and allowed to recover from 8:30-12:30 PM. All trials and behavior analyses were performed in the same way as above, with the addition of feeding (yes/no) as a behavior scored. This was used as a measure of tadpole normal behavior to determine if drugs had off-target effects, and tadpoles that did not eat were not included in the data set. The injector was blinded to the concentration and type of the drug being injected.

POMC is cleaved into three major peptides post-translationally, including aMSH, ACTH, and B-endorphin, which target different receptor families. To determine which peptide is relevant for tadpole aggressive behavior, we used pharmacology to manipulate each signaling pathway. A dose-response curve was performed for a selective MC4R antagonist, M4603 (Sigma Aldrich, #SHU9119), which binds aMSH. M4603 was injected icv at the dosages of 1.76 ug/uL (3 nmol), 0.589 ug/uL (1 nmol), and 0.1767 ug/uL (0.3 nmol) to determine the optimal concentration based on prior research using this drug in rainbow trout [53] (N = 5 per group; Figure S2). After determining the optimal concentration, tadpoles were injected with 1.76 ug/uL and control (saline) (aggression N = 10, control N = 10). ACTH was injected ip (0.05 ug/uL body mass; aggression N = 10, saline control N = 10). As the endorphin system targets many different opioid receptors, we tested the broad opioid receptor antagonist (naltrexone hydrochloride (Millipore-Sigma, #N3136)) injected ip at the dosage of 10 ug/uL (aggression N = 10, saline control N = 10). Due to the fast-acting nature of naltrexone [54,55], these tadpoles were injected between 7:30-8:30 AM and the aggression trials were run between 10:00 AM-12:00 PM.

### Data analysis

All statistics and figures were generated in R Studio (version 1.1.442) running R (version 3.5.2). To examine tadpole behavior, the glmmTMB R package [56] was used to run generalized linear mixed models. The linear mixed effect model was then followed with the Anova.glmmTMB function for reported statistical values.

Cell count data was examined using the glmmTMB R package to run generalized linear mixed models. For pS6, vasopressin, oxytocin, *pomc*, and colocalized cells, separate models were run using a negative binomial distribution appropriate for count data. For all models, we tested for significant differences between behavioral groups by including group, brain region, and their interaction as main effects. Tadpole identity was included as a random variable to account for repeated sampling of brain regions within individuals. Log of brain region area was included as an offset to account for potential differences in brain size. For colocalization data, the number of colocalized cells was included as the independent variable and the number of nonapeptide cells or *pomc* cells as a weight in the model. The model was then followed with the Anova.glmmTMB function for reported statistical values. When there was a significant interaction between group and brain region, a post-hoc test was run with emmeans R package (version 1.5.3) within brain regions and a Tukey-correction used for multiple hypothesis testing to adjust p-values for all post hoc comparisons.

PhosphoTRAP data was first processed by aligning trimmed reads to an *R.imitator* transcriptome and a count table was generated as previously described [50]. In order to determine differentially expressed genes from the phosphoTRAP analysis, a paired t-test was run comparing count values of each transcript between the INPUT and immunoprecipitated (IP) samples. This was done first within the “aggression” tadpoles, and second in the “control” tadpoles. Fold change was calculated of each transcript, which was averaged for each group and transformed using the log2 function in R (version 4.1.2). Differentially-expressed genes were identified based on a p-value below 0.05 and a fold change above 2.3. Finally, differentially-expressed gene lists were compared between aggression and control tadpoles. Transcripts that were differentially expressed in the same direction in both lists were removed, as those changes also seen in control animals were likely related to handling or stress. This generated a final list of transcripts with differential expression associated with aggression-related neuron activation (Supplementary Excel File).

## Results

### Aggressive behavior increases across development

Tadpole aggression increased across early development, including both the number of attacks (X2(5) = 40.234, p<0.001) and the attack duration (X2(5) = 75.376, p<0.001). The number of attacks increased after 4 weeks (p<0.001) while the duration of attacks increased in week 5 (p<0.001) (Figure 1).

**Figure 1.**
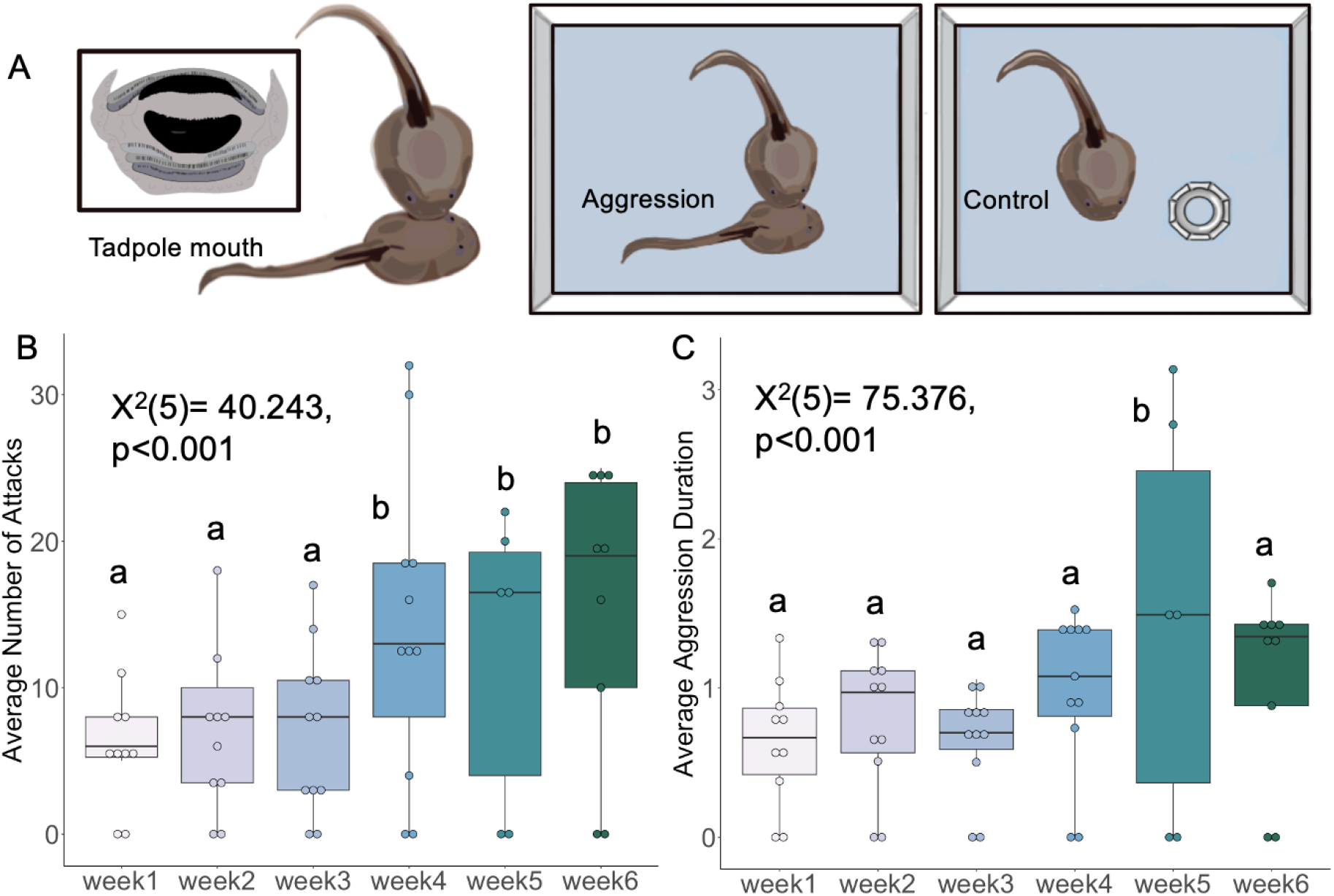
Tadpole aggression increases across development. **(A)** Schematic of behavioral assays run for aggressive behavior and controls. **(B)** The average number of attacks, quantified by the number of times the focal tadpole bit a conspecific, was significantly different between weeks 1-3 and weeks 4-6. **(C)** The average aggression duration, quantified by how long the focal tadpole aggressed, was significantly different between weeks 1-4 and week 5.

### Neural activity in the ventral hypothalamus increases in aggression

To identify brain regions that may be important in juvenile aggression, we compared general patterns of neural activity across several brain using immunohistochemical detection of phosphorylated ribosomes (pS6) and found differential activity across brain regions and behavioral groups (group*brain region: X2(6)=12.945, p=0.043; Figure 2). Post hoc analyses revealed that aggressive tadpoles have more pS6-positive cells in the ventral hypothalamus (t=3.109, p=0.002) compared to control animals.

**Figure 2.**
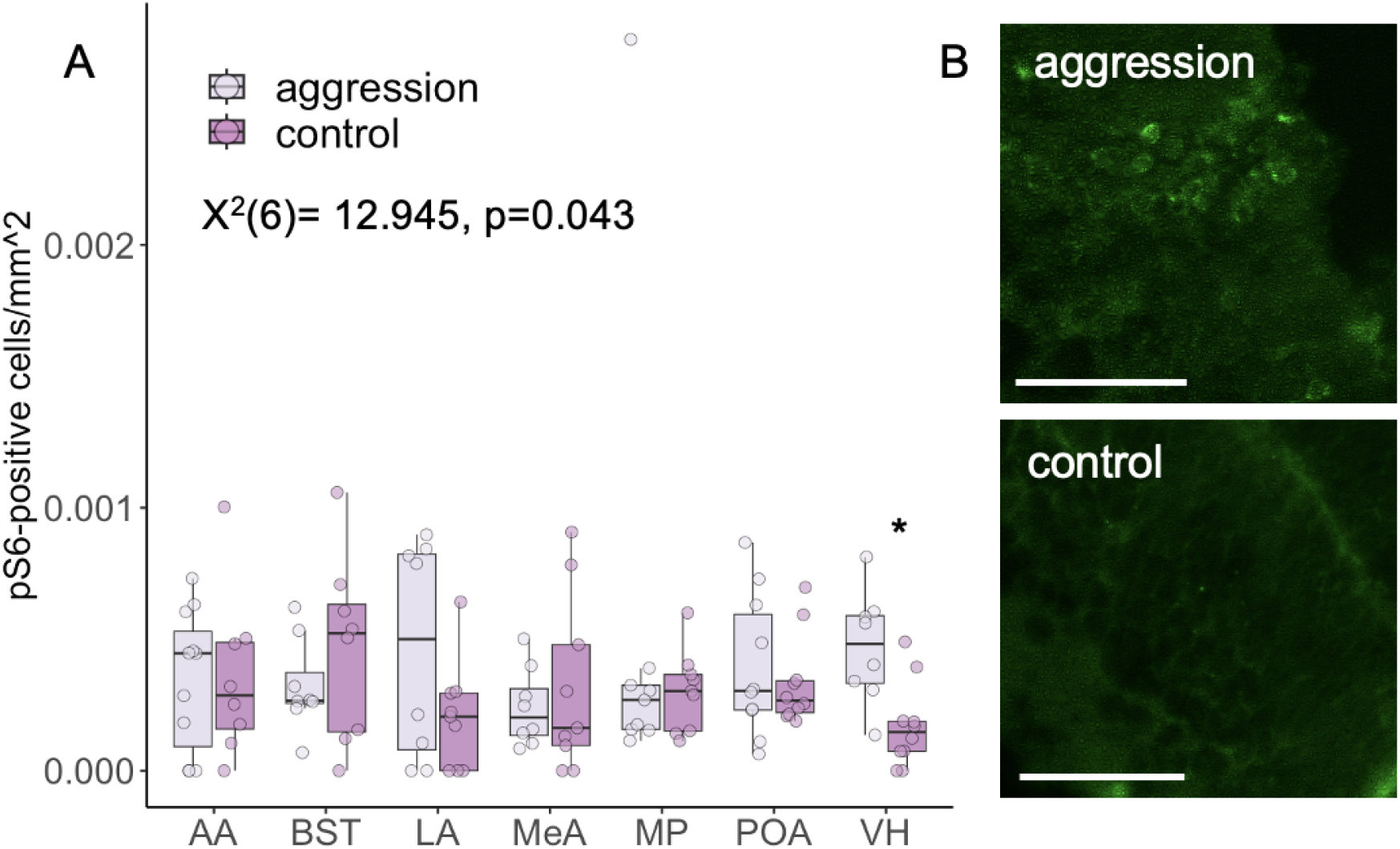
The ventral hypothalamus is more active in aggressive tadpoles. **(A)** pS6-positive cells were quantified across many brain regions to identify areas that may be important for juvenile aggression. The ventral hypothalamus shows a difference in activation between aggression and control animals (p = 0.03). **(B)** Micrographs of pS6-positive cells in the ventral hypothalamus of aggressive (top) and control (bottom) tadpoles; scale bar is 50um. Abbreviations: anterior amygdala (AA), bed nucleus of the stria terminalis (BST), lateral amygdala (LA), medial amygdala (MeA), medial pallium (MP), preoptic area of the hypothalamus (POA), ventral hypothalamus (VH).

Nonapeptides are involved in regulating juvenile play behavior in rodents [57] and were previously linked to tadpole aggression in a different poison frog species [18]. We examined the number and pS6 colocalization of oxytocin and vasopressin cells in aggression and control contexts (Figure S3). We found that the number of vasopressin cells in the preoptic area did not differ across groups (X2(1)=2.1861, p=0.13), and there was slight increase of colocalization of vasopressin with pS6 in control animals (X2(1)=3.346, p=0.067). We did not observe any differences in oxytocin cell number (X2(1)=0.8224, p=0.3645) or colocalization with pS6 (X2(1)=0.0013, p=0.9718) across groups.

### POMC neurons as a negative regulator of juvenile aggression through opioid receptors

We next took an untargeted approach to identify candidate cell types important for juvenile aggression using phosphoTRAP, a method for molecularly profiling transcripts being actively transcribed by cells [49] (Figure 3). In the telencephalon, there were 100 genes that were enriched and 325 genes that were depleted in immunoprecipitated RNA abundance compared to total RNA input (Figure S3A). In the diencephalon, there were 11 genes that were enriched and 932 genes that were depleted (Figure 3). In the spinal cord, there were 25 genes that were enhanced and 839 genes that were depleted (Figure S3C). We compared datasets from aggressive and control tadpoles, looking for signaling molecules or pathways that may be of importance in aggression and whose signaling pathways could be manipulated with pharmacology. We identified proopiomelanocortin (POMC), a precursor polypeptide with multiple post-translational peptides, as enriched (t=3.78, p=0.01) in the diencephalon phosphoTRAP dataset.

**Figure 3.**
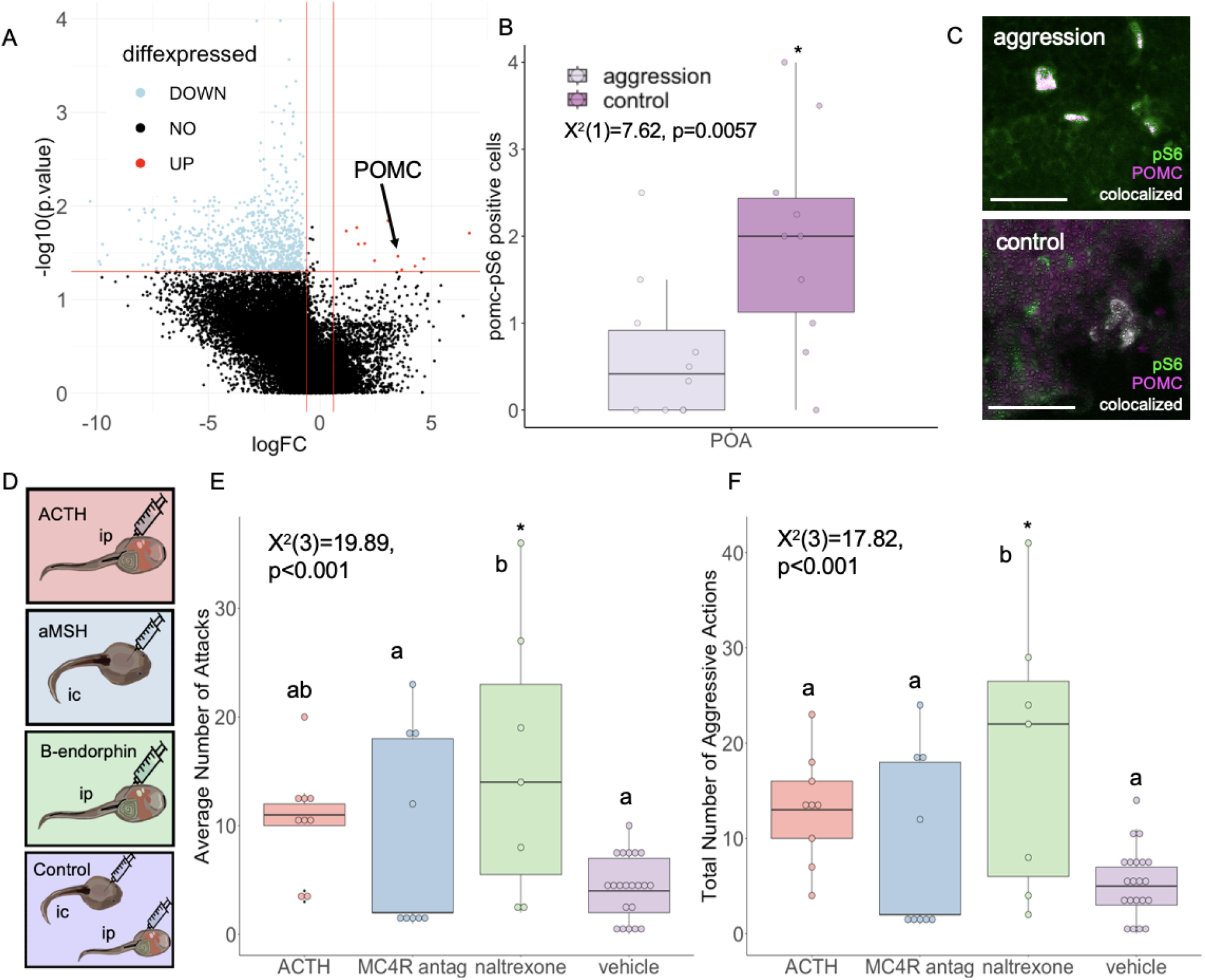
Proopiomelanocortin (POMC) negatively regulates aggression through opioid receptor signaling. **(A)** POMC is enriched in the diencephalon of aggressive tadpoles. Plot shows statistical significance plotted against magnitude of change. **(B)** There is a significant difference between POMC-pS6 positive cells in the preoptic area of the hypothalamus (POA) of aggressive and control animals. **(C)** Fluorescent imaging of POMC (magenta), pS6-positive cells (green) and their colocalization (white) in the POA; scale bar is 50 μm. **(D)** Schematic of experimental injection protocol. Treatment with naltrexone increased the number of attacks compared to MC4R antagonist and vehicle treated animals **(E)** and total aggressive actions compared to ACTH, MC4R antagonist, and vehicle treated animals **(F)**.

We next used fluorescent in situ hybridization to label *pomc* neurons and combined this with pS6 antibody staining (Figure 3B). There were very few *pomc-*positive cells in the arcuate nucleus or ventral hypothalamus at this stage in development (Figure S4). However, there were many *pomc*-positive cells in the preoptic area, where control animals had more *pomc* cells (X2(1)=7.622, p=0.006), Figure S5) and a higher colocalization of pS6 and *pomc* than aggressive animals (Figure 3B).

As POMC is post-translationally cleaved into different signaling peptides, we chose to pharmacologically manipulate these signaling pathways to determine which POMC-derived peptide was relevant to aggression in tadpoles (Figure 3D). We found a significant difference across drug treatments in the number of attacks (X2(3)=19.89, p<0.001) (Figure 3E) and total aggressive actions (X2(3)=17.82, p<0.001) (Figure 3F), where blocking opioid receptors increased number of bites (X2(1)=4.339, p<0.03) compared to controls. Manipulations with ACTH or blocking MC4R did not alter aggression compared to controls. This pharmacology data supports the histology data we collected, where decreased *pomc* signaling leads to increased aggression. Overall, our results suggest that the POMC-derived B-endorphin peptide is a negative regulator of aggressive behavior in tadpoles through opioid receptor signaling.

## Discussion

In an aggressive encounter, an animal must weigh the potential benefits of winning against the consequences of fighting, which may result in evolutionarily disadvantageous outcomes for the organism. Most aggression studies in vertebrate animals focus on adult males, with prominent roles of sex steroid-sensitive neurons in the ventromedial hypothalamus [2,8–13]. Although play fighting in laboratory juveniles suggests some mechanistic overlap, we still lack a basic understanding of juvenile aggression in the vertebrate brain. To address this knowledge gap, we investigated offensive aggression in *Ranitomeya imitator* tadpoles and found that aggressive behavior increases with age, is correlated with elevated neural activity in the ventral hypothalamus, and likely involves POMC acting as a negative regulator through opioid receptor signaling.

### Changes in aggressive behavior through development

In our study, we found that aggression increased over development. In many species, prior fighting experience increases aggression, including in rodents [58,59] and fish [60]. Winners of these outcomes will often win subsequent fights and this “winner effect” is mediated by androgens [61]. It is unclear from our study whether aggression increased with age due to repeated experience during aggression trials or if the brain regions or neuromodulators important for facilitating aggression develop after hatching. For example, it is possible that *pomc*-positive neurons in the preoptic area or μ opioid receptor distributions are still developing after tadpole hatching, as the developmental timing of *pomc*-neuron maturation in the hypothalamus and their signaling receptor distributions throughout the tadpole brain is unknown. Similarly, it may be possible that the connectivity of the ventral hypothalamus important for aggressive behavior develops after hatching. Examining the connectivity of the ventral hypothalamus as well as the distribution and abundance of *pomc* and opioid receptor neurons, coupled with independent tests of aggressive behavior, across development would clarify these remaining questions of behavioral development.

### The ventral hypothalamus is a conserved core brain region for aggression

In many adult mammals, the ventromedial hypothalamus, medial amygdala, and adjacent brain areas, also referred to as the “hypothalamic attack area”, are key regions for elicitation of aggression [45]. Here, we found increased pS6-positive cells in the ventral hypothalamus of tadpoles showing aggression compared to control animals presented with a novel object, consistent with studies in mammals. In a previous study in tadpoles of a closely related species, the Dyeing poison frog *Dendrobates tinctorius*, there was no increase in pS6-positive cells in animals that fought compared to those that did not in the ventral hypothalamus [18]. In that study, the medial pallium had increased pS6 in loser tadpoles compared to control animals that did not fight. However, that study used a different behavioral context (winners compared to losers and fighters compared to non-fighters), while our study compared animals that fought an intruder compared to those presented with a non-social object. More studies are needed to fully understand the role of the ventral hypothalamus in tadpole aggression moving forward. The present study is consistent with those in adult vertebrates that the ventral hypothalamus is a core brain region for governing aggression across development [2,8–13].

### Neuromodulators of aggression vary across development and species

Studies exploring aggression in adult mammals also support a role for many hormones and signaling molecules in regulating aggression [62]. For example, targeted gene expression profiling in zebrafish show multifactorial control of aggression, involving serotonin, somatostatin, dopamine, hypothalamo-neurohypophysial-system, hypothalamic-pituitary-interrenal, hypothalamo-pituitary-gonadal and histamine pathways [63]. The HPA axis has also been consistently linked to aggression, where low anxiety bred mice with higher circulating glucocorticoid levels have increased aggression compared to high anxiety strains [23]. Taken with our results, this suggests the HPA axis is a key node in a network of several brain regions that coordinate aggression in juvenile and adult animals.

The neural basis of aggression in vertebrate juveniles is most well understood in the context of “play fighting” in laboratory rodents, where vasopressin signaling plays a prominent role, ranging from juvenile play fighting to offensive aggression in adulthood [64]. Indeed, our previous study in *D. tinctorius* tadpoles suggested that nonapeptide cells (both vasopressin and oxytocin) are more active in tadpoles that win a fight compared to losers and control tadpoles that did not fight [18]. As such, we hypothesized that aggressive tadpoles would have more pS6-positive vasopressin or oxytocin cells and that one of these peptides would be enriched or depleted in our phosphoTRAP dataset. However, we did not find support for nonapeptide signaling being important for aggression in our present study, and we even found a trend for more vasopressin cells being pS6-positive in our control animals compared to the aggressive tadpoles. These contrasting findings could be due to different behavioral assays or species variation in the neural mechanisms of aggression. It should be noted that while many studies have investigated the effect of vasopressin on aggression in adults, results are variable. For example, when injecting adult mice with vasopressin in the lateral ventricle or amygdala there is no effect on aggression [65], whereas other studies in adult rats demonstrate a positive relationship between vasopressin injection in the anterior hypothalamus and aggression [48]. Put together, this indicates that vasopressin, although important in regulating adult aggression, may not be as important in regulating juvenile offensive aggression as was previously thought.

As nonapeptides and sex steroid hormones were not promising candidates for regulating tadpoles aggression in the present study, we next used an untargeted approach to identify transcripts enriched in pS6-positive neurons of aggressive tadpoles. We identified proopiomelanocortin (POMC), a precursor peptide with multiple post-translational peptides, as having increased translation by neurons activated during aggressive behavior. In humans, differential expression of POMC is linked to increased, abnormal aggressive displays in childhood and predisposes children to certain psychiatric disorders [45,66]. Further, differential methylation patterns of POMC are associated with abused children’s level of self-abuse ([67]. POMC is cleaved into three major peptides: aMSH, ACTH, and B-endorphin. To test the effects of each peptide on tadpole aggression, we pharmacologically manipulated each signaling pathway and found that blocking opioid receptors, which bind B-endorphins, lead to increased tadpole aggression. The post-translational modifications of POMC are complex and it is unclear why the *pomc* transcript was enriched in pS6-positive neurons during aggression and yet blocking opioid receptors increased aggression. Although our histology and pharmacology data was concordant, the phosphoTRAP data was not, as we would have expected depletion of *pomc* in aggressive tadpoles. Further studies into the specific regulation of these POMC peptide products would be required to fully delineate the relationships between aggression and each post-translational peptide.

B-endorphins and the opioid system underlie many behavioral and physiological effects. As B-endorphin binds the μ opioid receptor, this peptide has a well described role in analgesic effects, but is also seen to be involved in hedonics and homeostatic behaviors [68]. In particular, B-endorphins are proposed to modulate state switching in behavioral sequences [69], which is important for social behaviors and maintaining homeostasis [70]. In humans, certain psychiatric disorders characterized by abnormal aggression in childhood, such as borderline personality disorder, are thought to be a dysregulation of the endogenous opioid system [71]. Patients display symptoms such as self-injury, food restriction, aggression, and sensation seeking, which may be an attempt to stimulate the endogenous opioid system [72]. Alternatively, symptoms may result from alterations in the sensitivity of opioid receptors or availability of endogenous opioids after treatment with opioid receptor antagonists [73]. Individuals with autism spectrum disorder also display abnormal aggression, both towards others and themselves, that can be ameliorated by manipulating the POMC system. When administered an opioid blocker, autistic patients exhibited reduced self-injurious behavior [74]. Studies in animals offer conflicting conclusions, where in some studies endorphins decrease aggression and B-endorphin levels were negatively correlated with aggressive actions [75], while in another study naltrexone increases aggressive behaviors in male mice [76], similar to our finding in tadpoles. More studies in adults and juveniles are needed to determine if the opioid system is a conserved mediator of aggression across development and species.

Although we did not observe an effect through manipulation of aMSH or ACTH in tadpole aggression, studies in other taxa have demonstrated roles for both these peptides. For example, in cichlid fish, treatment with exogenous aMSH increases aggression in male adults [77], and injecting female rats with aMSH also increases aggression [78]. Importantly, aMSH produced by POMC cleavage in the arcuate nucleus regulates food intake in many taxa, where aMSH binding to MC4R receptors suppresses appetite [79]. We did not observe an effect of aMSH treatment on *R. imitator* tadpoles, which reinforces findings in *O. pumilio* tadpoles that aggression and hunger are decoupled [44]. Aggression studies with ACTH also suggest a role for promoting aggression, where mice injected with ACTH attack intruders more quickly [80], and rats bred for low anxiety had higher plasma corticotropin (ACTH) and subsequently exhibited higher levels of aggression as compared to high anxiety-bred or control rats [81]. Interestingly, injecting mice daily with ACTH will initially lead to an increase in aggression, but causes a decrease in aggressive action by days 4-7 [82]. Most of these studies of the influence of aMSH and ACTH on aggression is conducted in males and thus it is likely that age can have varying effects on behavior.

### Summary

Our results suggest that POMC is a negative regulator of tadpole aggression through opioid receptor signaling. The neural distribution of opioid receptors in amphibians is unknown and future work will focus on where these receptors are located, which opioid receptors are important for tadpoles aggression, and how these opioid systems may be different in aggressive and non-aggressive tadpole species. More broadly, this work provides a foundation for examining underlying neural mechanisms of juvenile aggression in vertebrates.

## Acknowledgements

The authors acknowledge that this research was conducted on the ancestral lands of the Muwekma Ohlone people at Stanford. We understand the implications of the historical and present colonialism these people experience and celebrate their continued stewardship of their lands. We would also like to thank the Laboratory of Organismal Biology for their assistance with animal care, helpful discussions, and guidance throughout this project.

## Data accessibility

All raw data, analysis scripts, and intermediate data analysis files will be publicly available at the time of publication. Depending on the file type, these will be available through NCBI (BioProjects/SRA) or DataDryad.

