## Supplementary Data for "Proopiomelanocortin (POMC) is a negative regulator of tadpole aggression through opioid receptor signaling"

### Supplementary Materials

| WEEK | Aggression mean | Aggression sd | Stimuli mean | Stimuli sd |
|---|---|---|---|---|
| 1 | 14.32 mm | 1.647mm | 13.00mm | 0.14mm |
| 2 | 17.90mm | 1.746mm | 16.35mm | 1.34mm |
| 3 | 20.37mm | 1.62mm | 19.15mm | 0.77mm |
| 4 | 21.42mm | 1.16mm | 20.55mm | 0.21mm |
| 5 | 22.15mm | 1.31mm | 21.50mm | 1.55mm |
| 6 | 22.92mm | 1.89mm | 23.60mm | 1.41mm |

**Table S1. Longitudinal study tadpole size. (A)** Table of average sizes and standard deviations of aggression and stimuli tadpoles over 6 weeks.

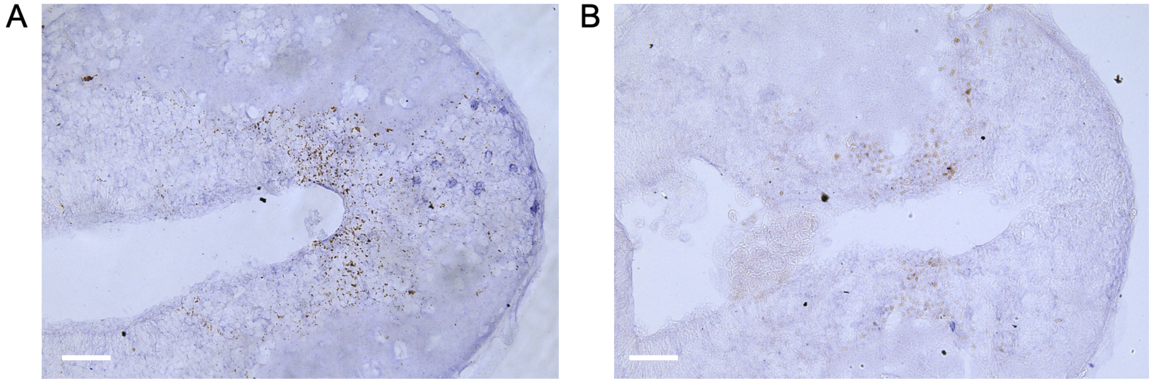


**Figure S1. Antisense and sense *pomc* probe test. (A)** Image of *pomc* antisense probe; scale bar = 50um. **(B)** Image of *pomc* sense probe; scale bar = 50um.

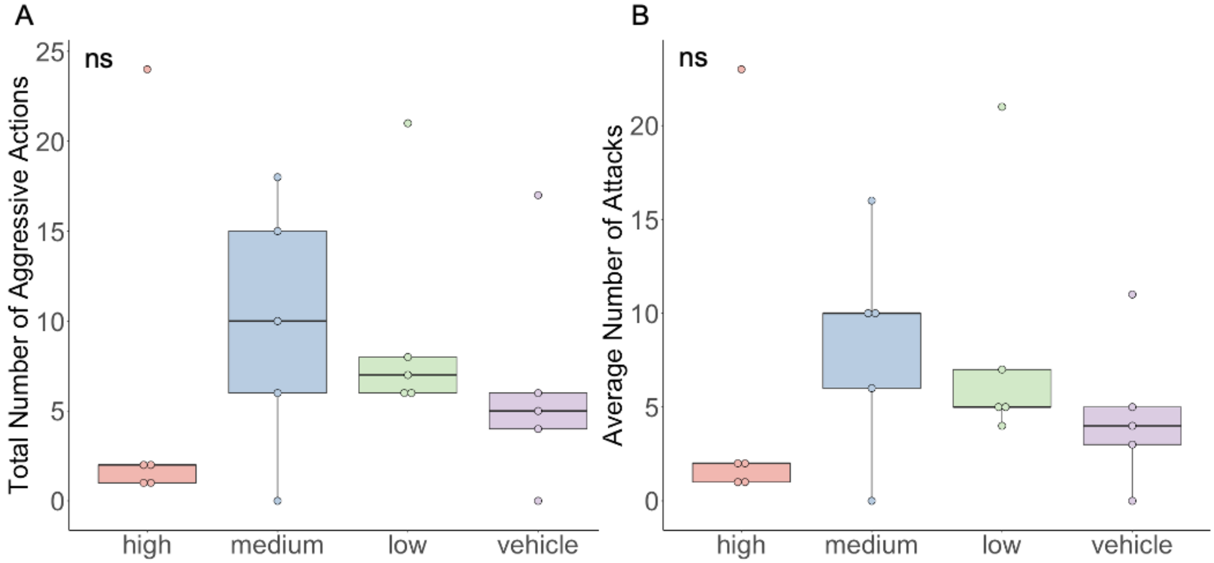


**Figure S2. aMSH antagonist dose-response trial. (A)** Average number of attacks between different dosages of an aMSH antagonist. **(B)** Total number of aggressive actions (bite, chase, thrash) between different dosages of an aMSH antagonist.

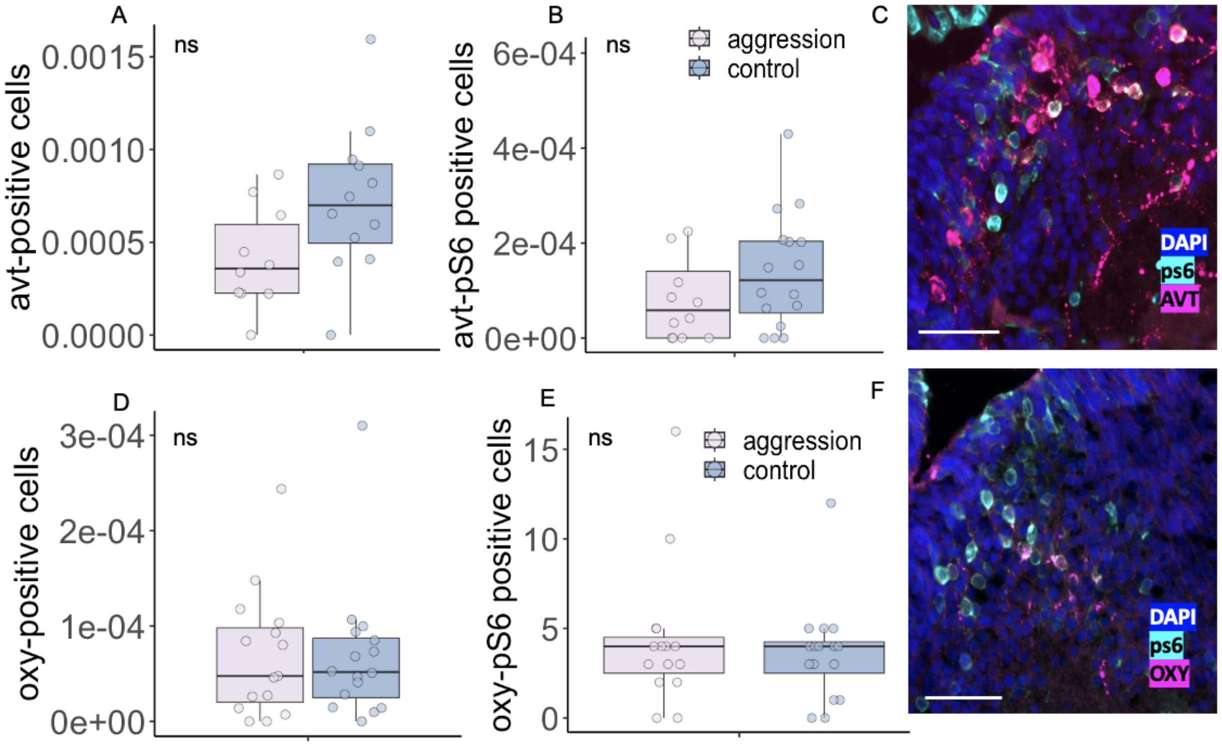


**Figure S3. Nonapeptide cell number and pS6 colocalization do not differ between aggression and control animals. (A)** The number of AVT cells in aggression and control animals is not significantly different. **(B)** The number of AVT-ps6 positive cells in aggression and control animals is not different. **(C)** Fluorescent imaging of AVT (magenta)-pS6 (cyan) positive cells in the medial preoptic area; scale bar= 50µm. **(D)** The number of oxytocin positive cells is not different between aggression and control animals. **(E)** The number of oxytocin (magenta)-pS6 (cyan) positive cells is not different between aggression and control animals. **(F)** Fluorescent imaging of OXY-ps6 positive cells in the medial preoptic area in an aggressive tadpole; scale bar= 50µm.

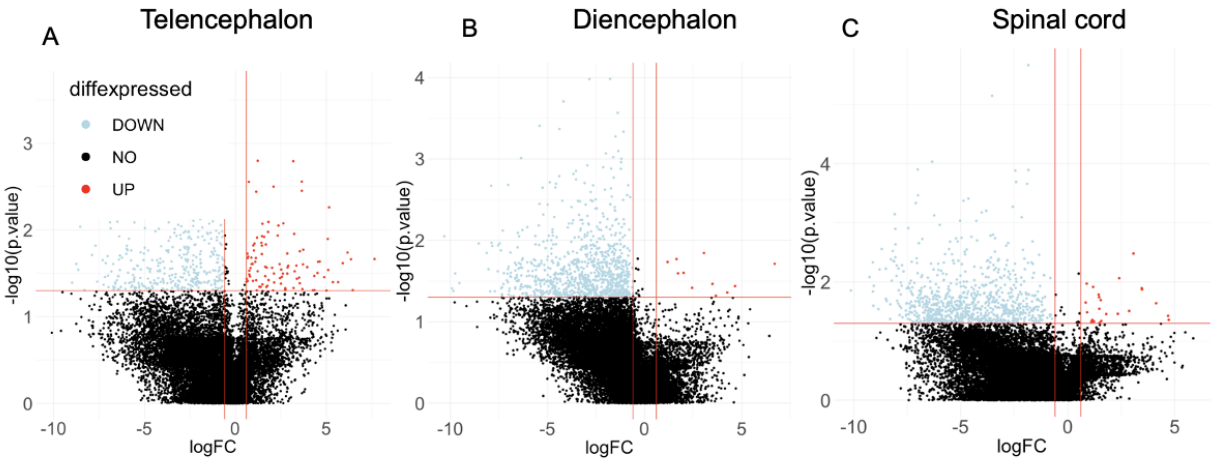


**Figure S3. Molecular profiling of active neurons using PhosphoTRAP. (A)** Molecular profiling of behaviorally-relevant genes in the telencephalon. Plot shows statistical significance plotted against magnitude of change. **(B)** Molecular profiling of behaviorally-relevant genes in the diencephalon. **(C)** Molecular profiling of behaviorally-relevant genes in the spinal cord. Key: light blue, depleted transcripts; red, enriched transcripts; black, transcripts not significantly enriched or depleted.

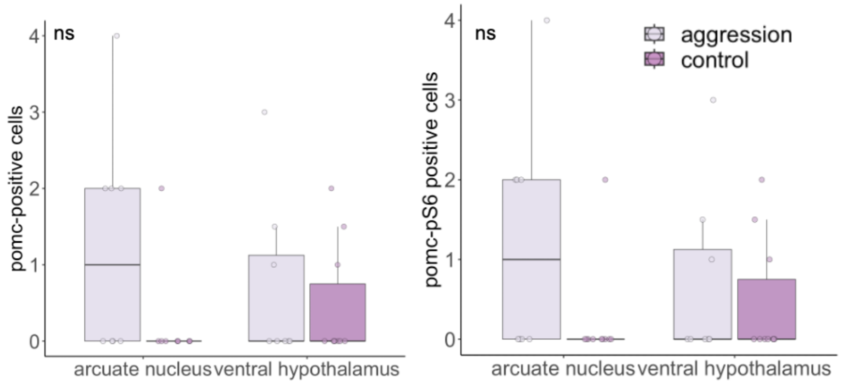


**Figure S4. POMC cell in situ hybridization cell counts. (A)** There was not a significant difference between POMC cell numbers in the arcuate nucleus and ventral hypothalamus. **(B)** There was not a significant difference between POMC-pS6 colocalization in the arcuate nucleus and ventral hypothalamus.

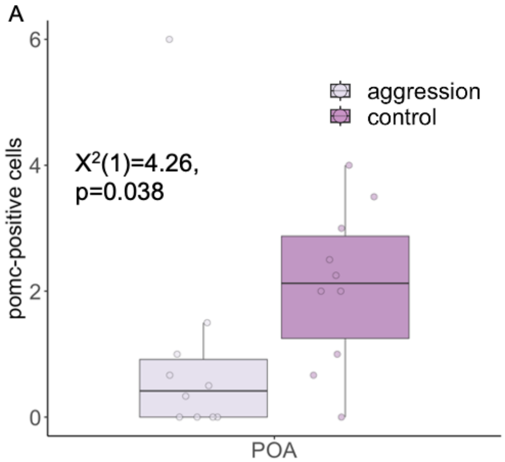


**Figure S5. POMC cell in situ hybridization. (A)** There was a significant difference between POMC cell numbers in the preoptic area (POA).
